# A novel lineage of candidate pheromone receptors for sex communication in moths

**DOI:** 10.1101/707174

**Authors:** Lucie Bastin-Héline, Arthur de Fouchier, Song Cao, Fotini Koutroumpa, Gabriela Caballero-Vidal, Stefania Robakiewicz, Christelle Monsempes, Marie-Christine François, Tatiana Ribeyre, Anne de Cian, William B. Walker, Guirong Wang, Emmanuelle Jacquin-Joly, Nicolas Montagné

**Affiliations:** INRA-CAAS International Joint Laboratory for Plant Protection “Bacteria-Insect-Plant Interaction for pest biocontrol”; Sorbonne Université, Inra, CNRS, IRD, UPEC, Université Paris Diderot, Institute of Ecology and Environmental Sciences of Paris, Paris and Versailles, France; State Key Laboratory for Biology of Plant Diseases and Insect Pests, Institute of Plant Protection, Chinese Academy of Agricultural Sciences, Beijing, China; CNRS UMR 7196, INSERM U1154, Museum National d’Histoire Naturelle, Paris, France; Department of Plant Protection Biology, Swedish University of Agricultural Sciences, Alnarp, Sweden

## Abstract

Sex pheromone receptors (PRs) are key players in chemical communication between mating partners in insects. In the highly diversified insect order Lepidoptera, male PRs tuned to female-emitted type I pheromones (which make up the vast majority of pheromones identified) form a dedicated subfamily of odorant receptors (ORs). Here, using a combination of heterologous expression and *in vivo* genome editing methods, we bring functional evidence that at least one moth PR does not belong to this subfamily but to a distantly related OR lineage. This PR, identified in the cotton leafworm *Spodoptera littoralis*, is over-expressed in male antennae and is specifically tuned to the major sex pheromone component emitted by females. Together with a comprehensive phylogenetic analysis of moth ORs, our functional data suggest two independent apparitions of PRs tuned to type I pheromones in Lepidoptera, opening up a new path for studying the evolution of moth pheromone communication.

## Introduction

The use of pheromone signals for mate recognition is widespread in animals, and changes in sex pheromone communication are expected to play a major role in the rise of reproductive barriers and the emergence of new species (Smadja and Butlin, 2009). Since the first chemical identification of such a pheromone in *Bombyx mori* (Butenandt et al., 1961), moths (Insecta, Lepidoptera) have been a preferred taxon for pheromone research (Cardé and Haynes, 2004; Kaissling, 2014). The diversification of pheromone signals has likely played a prominent role in the extensive radiation observed in Lepidoptera, which represents almost 10% of the total described species of living organisms (Stork, 2018).

Female moths generally release a species-specific blend of volatile molecules, which attract males over a long distance (Cardé and Haynes, 2004). Four types of sex pheromones have been described in moths (types 0, I, II and III), with distinct chemical structures and biosynthetic pathways (Löfstedt et al., 2016). 75% of all known moth sex pheromone compounds belong to type I and are straight-chain acetates, alcohols or aldehydes with 10 to 18 carbon atoms (Ando et al., 2004). Type I pheromones have been found in most moth families investigated, whereas the other types are restricted to only a few families (Löfstedt et al., 2016).

In moth male antennae, pheromone compounds are detected by dedicated populations of olfactory sensory neurons (OSNs). Each type of OSN usually expresses one pheromone receptor (PR) responsible for signal transduction. PRs are 7-transmembrane domain proteins belonging to the odorant receptor (OR) family and, as ORs, are co-expressed in OSNs together with the conserved co-receptor ORco (Chertemps, 2017; Fleischer and Krieger, 2018).

Since the first discovery of moth PRs (Krieger et al., 2004; Sakurai et al., 2004), numerous pheromone receptors tuned to type I pheromone compounds have been characterized through different hererologous expression systems, and most appeared to be specific to only one compound (Zhang and Löfstedt, 2015). More recently, a few receptors tuned to type 0 and type II pheromones have also been characterized (Zhang et al., 2016; Yuvaraj et al., 2017). Type I PRs belong to a dedicated monophyletic subfamily of ORs, the so-called “PR clade”, suggesting a unique emergence early in the evolution of Lepidoptera (Yuvaraj et al., 2018). Another hallmark of type I PRs is their male-biased expression (Koenig et al., 2015). The phylogenetic position and the expression pattern have thus been the main criteria used to select candidate PRs for functional studies.

Likewise, we selected PRs from the male transcriptome of the cotton leafworm *Spodoptera littoralis* (Legeai et al., 2011), a polyphagous crop pest that uses type I sex pheromone compounds (Muñoz et al., 2008) and that has been established as a model in insect chemical ecology (Ljungberg et al., 1993; Binyameen et al., 2012; Saveer et al., 2012; Poivet et al., 2012; de Fouchier et al., 2017). Through heterologous expression, we characterized two PRs tuned to minor components of the *S. littoralis* pheromone blend (Montagné et al., 2012; de Fouchier et al., 2015), but none of the tested candidate PRs detected the major pheromone component (*Z,E*)-9,11-tetradecadienyl acetate (hereafter referred as (*Z,E*)-9,11-14:OAc), which is necessary and sufficient to elicit all steps of the male mate-seeking behavioral sequence (Quero et al., 1996).

In order to identify new type I PR candidates, we focused on male-biased ORs, whether they belong to the PR clade or not. Notably, a preliminary analysis of *S. littoralis* OR expression patterns led to the identification of such a receptor, SlitOR5, which was over-expressed in male antennae but did not belong to the PR clade (Legeai et al., 2011). Furthermore, a recent RNAseq analysis showed that SlitOR5 was the most abundant OR in *S. littoralis* male antennae (Walker et al., 2019). Here, we demonstrate that SlitOR5 is the receptor for (*Z,E*)-9,11-14:OAc using a combination of heterologous expression and *in vivo* genome editing methods. Based on a comprehensive phylogenetic analysis of lepidopteran ORs, we show that SlitOR5 belongs to an OR subfamily that is distantly related to the PR clade but harbors numerous sex-biased ORs from distinct moth families. Altogether, these results suggest that PRs detecting type I pheromones evolved at least twice in Lepidoptera, which offers a more detailed and complex panorama on moth PR evolution.

## Results

### SlitOr5 is over-expressed in males but does not belong to the type I pheromone receptor clade

We first used quantitative real-time PCR to compare the relative expression levels of the *SlitOr5* gene in *S. littoralis* male and female adult antennae. We found *SlitOr5* expressed with a more than fifty-fold enrichment in the male antennae (Fig. 1A), thus confirming previous observations (Legeai et al., 2011; Walker et al., 2019).

**Figure 1.**
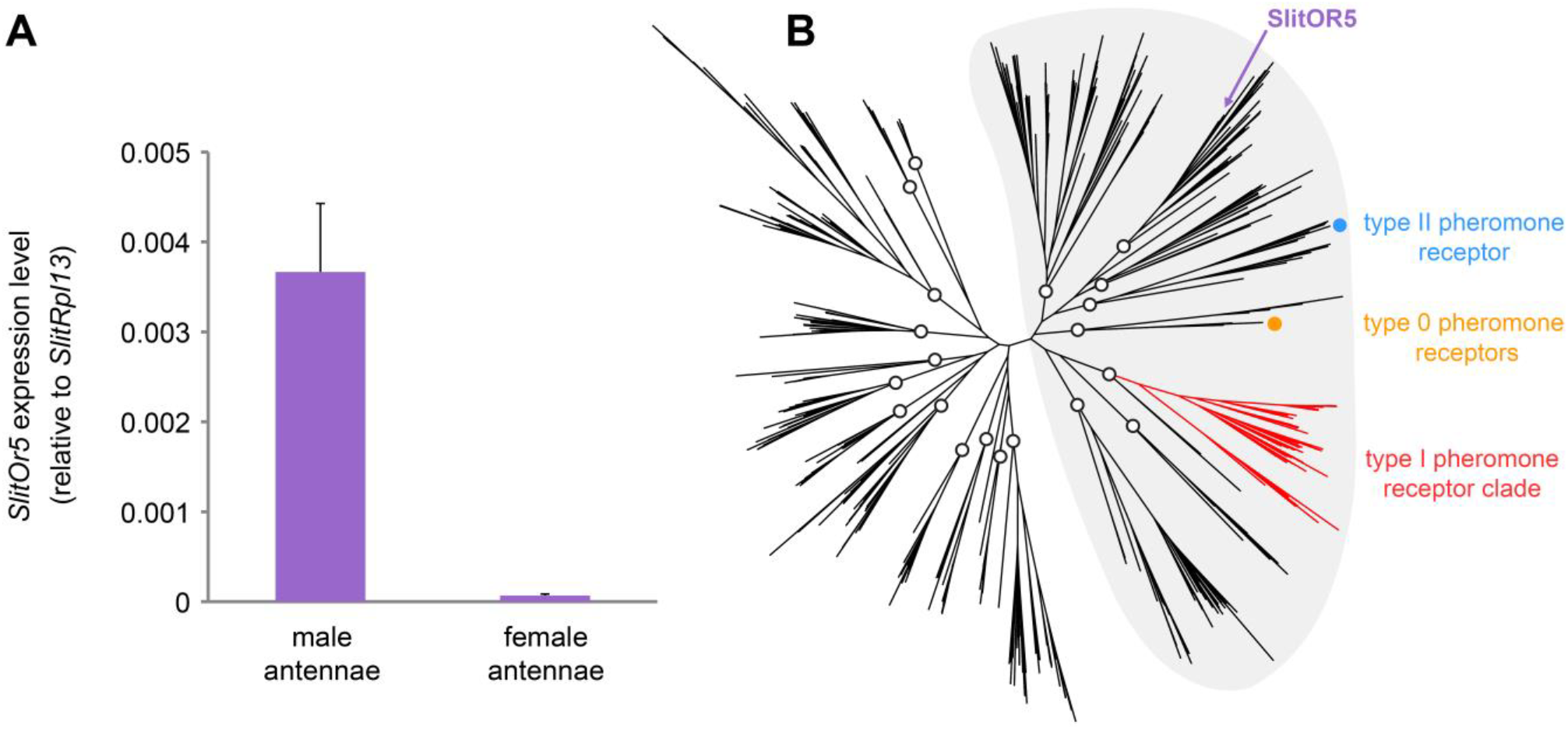
*SlitOr5* is over-expressed in males but does not belong to the pheromone receptor clade. **(A)** Expression levels of *SlitOr5* in adult male and female antennae of *S. littoralis*, as measured by real-time qPCR. Expression levels have been normalized to the expression of *SlitRpl13*. Plotted values represent the mean normalized expression values ± SEM (*n* = 3). Raw results are available in Figure 1 – source data 1. **(B)** Unrooted maximum likelihood phylogeny of lepidopteran ORs, based on 506 amino acid sequences from 9 species, each belonging to a different superfamily. The position of SlitOR5 and of receptors for type 0, type I and type II pheromone compounds is highlighted. Circles on the nodes indicate the distinct paralogous OR lineages, supported by a transfer bootstrap expectation (TBE) > 0.9. All the PR-containing lineages grouped within a large clade (highlighted in grey) also supported by the bootstrap analysis. The sequence alignment file is available in Figure 1 – source data 2.

We reconstructed a maximum likelihood phylogeny of lepidopteran ORs, based on entire OR repertoires from 9 different species. Among the 20 paralogous lineages identified (each having evolved in principle from an ancestral OR present in the last common ancestor of Lepidoptera), SlitOR5 belonged to a lineage distantly related to the type I PR clade, as well as to the lineages containing type 0 and type II PRs (Fig. 1B). These four paralogous lineages grouped within a larger clade highly supported by the bootstrap analysis (highlighted in grey in Fig. 1B). This clade has been previously shown to contain ORs tuned to terpenes and aliphatic molecules – including sex pheromones – and exhibits higher evolutionary rates compared to more ancient clades that contain many receptors for aromatics (de Fouchier et al., 2017).

### SlitOR5 binds (Z,E)-9,11-14:OAc with high specificity and sensitivity

We next used two complementary heterologous systems to characterize the function of SlitOR5 and assess whether it is the receptor to (*Z,E*)-9,11-14:OAc, the major component of the *S. littoralis* sex pheromone blend. First, we expressed SlitOR5 in *Drosophila melanogaster* OSNs housed in at1 trichoid sensilla, in place of the endogenous PR DmelOR67d (Kurtovic et al., 2007). Single-sensillum recordings were performed to measure the response of at1 OSNs to 26 type I pheromone compounds (Supplementary File 1), including all the components identified in the pheromones of *Spodoptera* species (El Sayed, 2018). SlitOR5-expressing OSNs strongly responded to (*Z,E*)-9,11-14:OAc (65 ±15 spikes.s^−1^), whereas there was no significant response to any other compound (Fig. 2A).

**Figure 2.**
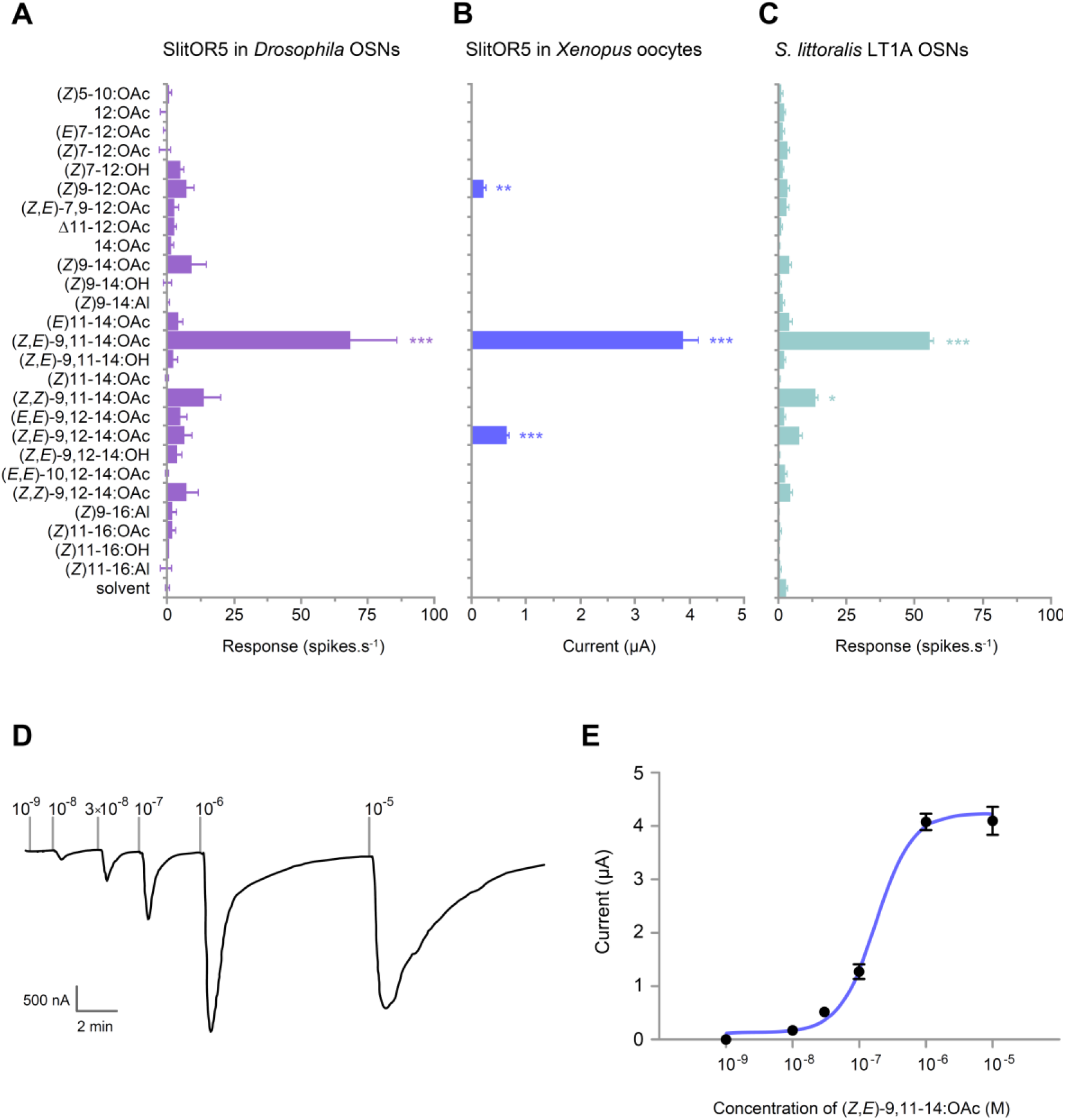
SlitOR5 is the receptor for the major component of the *S. littoralis* pheromone blend. **(A)** Action potential frequency of *Drosophila* at1 OSNs expressing SlitOR5 (*n* = 8) after stimulation with 26 type I pheromone compounds (10 µg loaded in the stimulus cartridge). *** *P* < 0.001, significantly different from the response to solvent (one-way ANOVA followed by a Tukey’s *post hoc* test). **(B)** Inward current measured in *Xenopus* oocytes co-expressing SlitOR5 and SlitORco (*n* = 13-16) after stimulation with the same panel of pheromone compounds (10^−4^ M solution). *** *P* < 0.001, ** *P* < 0.01, significantly different from 0 (Wilcoxon signed rank test). **(C)** Action potential frequency of LT1A OSNs from *S. littoralis* male antennae (*n* = 8-16) after stimulation with pheromone compounds (1 µg loaded in the stimulus cartridge). *** *P* < 0.001, * *P* < 0.1, significantly different from the response to solvent (one-way ANOVA followed by a Tukey’s *post hoc* test). **(D)** Representative trace showing the response of a *Xenopus* oocyte co-expressing SlitOR5 and SlitORco after stimulation with a range of (*Z,E*)-9,11-14:OAc doses from 10^−9^ M to 10^−5^ M. **(E)** Dose-response curve of SlitOR5/ORco *Xenopus* oocytes (*n* = 9) stimulated with (*Z,E*)-9,11-14:OAc (EC_50_ = 1.707×10^-7^ M). Plotted values in (A-C and E) are mean responses ± SEM. Raw results for all experiments are available in Figure 2 – source data 1.

Then, we co-expressed SlitOR5 with its co-receptor SlitOrco in *Xenopus* oocytes and recorded the responses to the same panel of pheromone compounds using two-electrode voltage-clamp. A strong current was induced in SlitOR5-expressing oocytes when stimulated with (*Z,E*)-9,11-14:OAc (3.9 ±0.3 µA), whereas only minor currents were recorded in response to (*Z,E*)-9,12-14:OAc and (*Z*)9-12:OAc (Fig. 2B). SlitOR5 sensitivity was assessed with a dose-response experiment that showed a low detection threshold (10^−8^ M) and an EC_50_ of 1.707×10^−7^ M (Fig. 2D-E).

We compared the response spectra of heterologously expressed SlitOR5 with that of *S. littoralis* male OSNs housed in type 1 long trichoid sensilla (LT1A OSNs, Fig. 2C), known to detect (*Z,E*)-9,11-14:OAc (Ljungberg et al., 1993; Quero et al., 1996). When stimulated with the 26 pheromone compounds, LT1A OSNs significantly responded to (*Z,E*)-9,11-14:OAc (55 ±4 spikes.s^−1^) and to a lesser extent to its stereoisomer (*Z,Z*)-9,11-14:OAc, which is absent from any *Spodoptera* pheromone. This mirrored the response spectra of heterologously expressed SlitOR5, especially the one observed in *Drosophila* OSNs (Fig. 2A).

### In vivo response to (Z,E)-9,11-14:OAc is abolished in SlitOr5 mutant males

To confirm *in vivo* that SlitOR5 is the receptor for the major sex pheromone component of *S. littoralis*, we carried out a loss-of-function study by generating mutant insects for the gene *SlitOr5*. The CRISPR/Cas9 genome editing system was used to create a mutation in the first exon of *SlitOr5* with the aim of disrupting the open reading frame. Guide RNAs were injected along with the Cas9 protein in more than one thousand eggs. We genotyped 66 G0 hatched larvae and found 7 individuals carrying at least one mutation in *SlitOr5*. These 7 G0 individuals were back-crossed with wild-type individuals to create 7 G1 heterozygous mutant lines. We selected a line carrying a single mutation that consisted of a 10-bp deletion at the expected location, introducing a premature STOP codon within the *SlitOr5* transcript after 247 codons (Fig. 3A).

**Figure 3.**
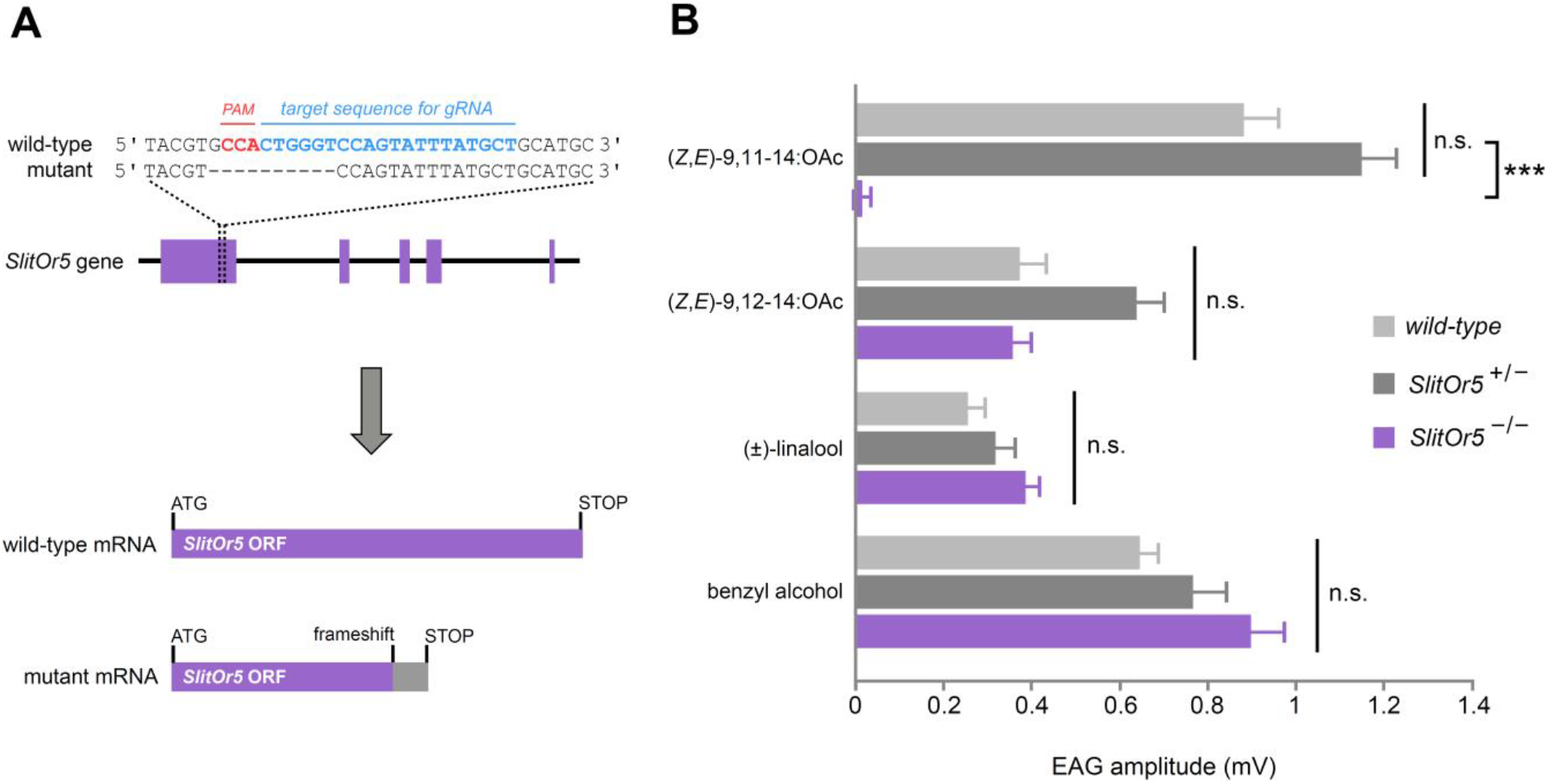
Response to the major pheromone component is abolished in *SlitOr5* mutants. **(A)** Location of the 10-bp deletion induced in the first exon of the *SlitOr5* gene by the CRISPR/Cas9 system. The sequence complementary to the RNA guide is indicated in blue, and the protospacer adjacent motif (PAM) in red. The frameshift created in the *SlitOr5* open reading frame (ORF) induced a premature stop codon. **(B)** Electroantennogram (EAG) amplitude measured in *S. littoralis* male antennae isolated from wild-type animals (light grey, *n* = 14), heterozygous *SlitOr5* mutants (dark grey, *n* = 18) and homozygous *SlitOr5* mutants (purple, *n* = 8) after stimulation with pheromone compounds (1 µg in the stimulus cartridge) and plant volatiles (10 µg in the stimulus cartridge). Plotted values represent the normalized mean response ± SEM (response to the solvent was subtracted). *** *P* < 0.001, significantly different from the response of the other genotypes; n.s.: not significantly different (one-way ANOVA, followed by a Tukey’s *post hoc* test). Raw results for the EAG experiment are available in Figure 3 – source data 1.

We next generated G2 homozygous mutant males (*SlitOr5*^−/-^) and compared their ability to detect (*Z,E*)-9,11-14:OAc to that of wild-type and heterozygous (*SlitOr5*^+/−^) males, using electroantennogram (EAG) recordings (Fig. 3B). When stimulated with (*Z,E*)-9,11-14:OAc, wild-type and *SlitOr5*^+/−^ antennae exhibited similar EAG amplitudes (0.89 mV and 1.16 mV, respectively), whereas the response was completely abolished in *SlitOr5*^−/−^ antennae (0.02 mV). Control experiments using a *S. littoralis* minor pheromone component and plant volatiles known to induce EAG responses in *S. littoralis* (Saveer et al., 2012; López et al., 2017) showed that antennal responses were not impaired in *SlitOr5*^−/−^ mutants, as these odorants elicited similar responses in wild-type, heterozygous and homozygous moths (Fig. 3B). When challenged with wild-type virgin mature females during the scotophase, none of the homozygous male mutants (*n* = 10) exhibited classical wing fanning and clasper extrusion as did wild-type males, and they did not mate over several days of observation. Homozygous female mutants (*n* = 10), however, all did mate with wild-type virgin males, usually within one day. Overall, these results confirm that SlitOR5 is the receptor responsible for the detection of the major component of the *S. littoralis* female pheromone blend.

### A novel lineage of candidate moth pheromone receptors

In view of these results and the unexpected phylogenetic position of SlitOR5, we rebuilt the phylogeny of the lepidopteran OR clade containing SlitOR5 and the known receptors for type 0, type I and type II pheromones (highlighted in grey in Fig. 1B), adding all ORs showing a strong sex-biased expression (at least 10-fold in one sex compared to the other) and ORs whose ligands were known as of September 2018 (Supplementary File 2). ORs grouped within 8 different paralogous lineages, four of which including PRs (Fig. 4). One was the so-called PR clade that, as previously observed, contained all type I PRs characterized so far (except SlitOR5) as well as two type II PRs (Zhang et al., 2016). The other three lineages harboring PRs consisted of one containing SlitOR5, one containing EgriOR31 (a type II PR from the geometrid *Ectropis grisescens*; Li et al., 2017) and one containing EsemOR3 and 5 (type 0 PRs from the non-dytrisian moth *Eriocrania semipurpurella*; Yuvaraj et al., 2017). Interestingly, most sex-biased lepidopteran ORs identified to date clustered within the PR clade and the lineage containing SlitOR5. While sex-biased ORs within the PR clade were mainly male-biased, the SlitOR5 lineage contained an equal proportion of male and female-biased receptors, identified from species belonging to different families of Lepidoptera. Deep nodes within the phylogeny were highly supported by the bootstrap analysis, enabling us to state that these two PR-containing clades were more closely related to clades harboring receptors for plant volatiles than to each other. This suggests that receptors tuned to type I pheromone compounds emerged twice independently during the evolution of Lepidoptera, and that the clade containing SlitOR5 may constitute a novel lineage of candidate PRs.

**Figure 4.**
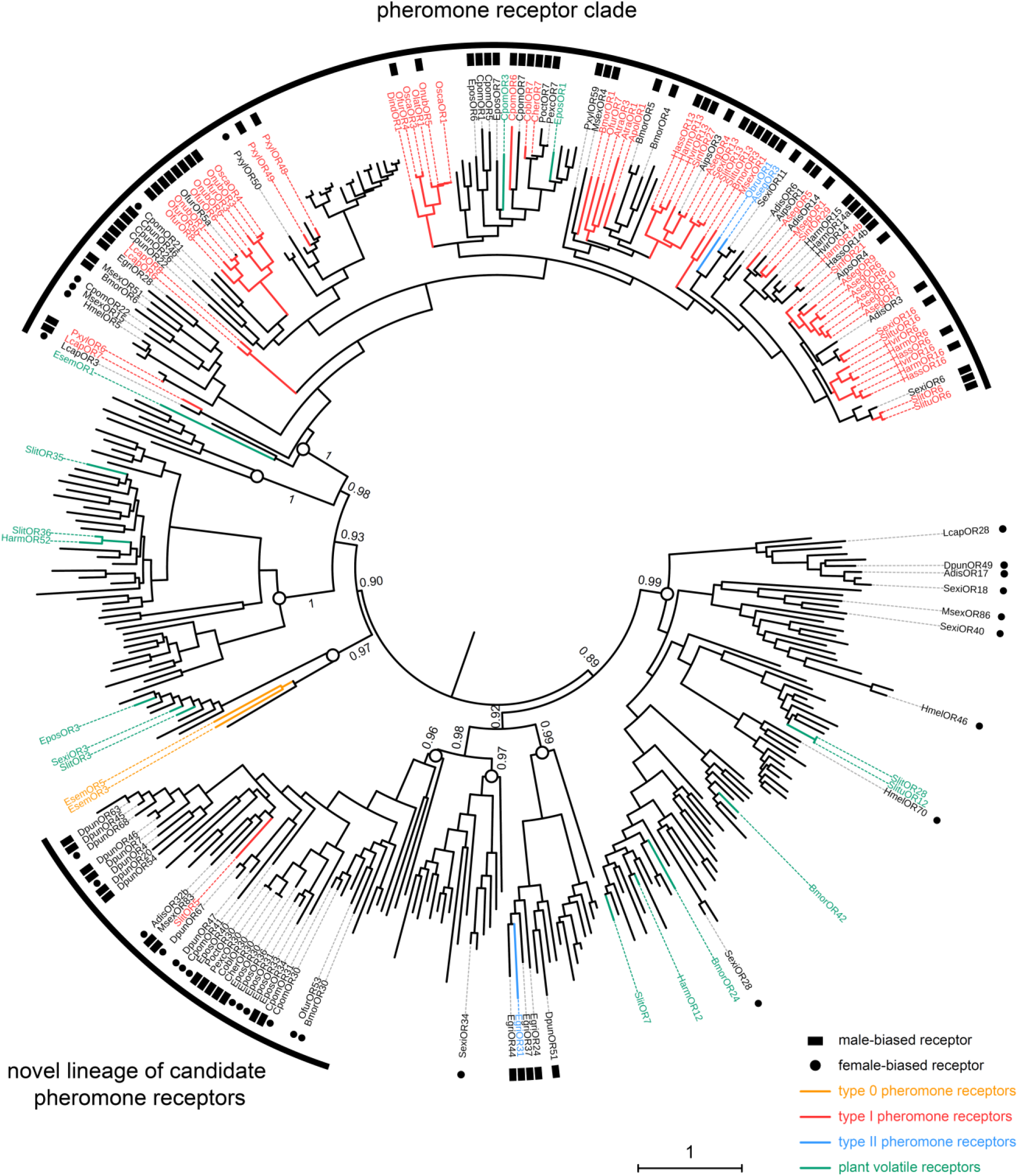
SlitOR5 may define a novel lineage of candidate pheromone receptors in Lepidoptera. Maximum likelihood phylogeny of the lepidopteran OR clade that includes all the paralogous lineages containing pheromone receptors. 360 sequences from 34 lepidopteran species were included. Functional and expression data shown on the figure have been compiled from the literature (Supplementary File 2). Branch colors indicate OR function, when characterized: PRs for type I pheromones are depicted in red, those for type II pheromones in blue and those for type 0 pheromones in orange. ORs tuned to plant volatiles are depicted in green. Symbols at the edge indicate expression data: male-biased ORs are highlighted with black squares and female-biased ORs with black dots. Circles on the nodes indicate the distinct paralogous OR lineages. Support values on basal nodes are transfer bootstrap expectation (TBE) values. The tree has been rooted using an outgroup as identified in the lepidopteran OR phylogeny shown in Fig. 1. The scale bar indicates the expected number of amino acid substitutions per site. The sequence alignment file is available in Figure 4 – source data 1.

## Discussion

While moth sex pheromone receptors have been the most investigated ORs in Lepidoptera, with more than sixty being functionally characterized (Zhang and Löfstedt, 2015), it remains unclear how and when these specialized receptors arose. Type I PRs have been proposed to form a monophyletic, specialized clade of ORs, the so-called “PR clade”, which emerged early in the evolution of Lepidoptera (Yuvaraj et al., 2017, 2018). Here, we bring functional and phylogenetic evidence that type I PRs are not restricted to this clade and likely appeared twice independently in Lepidoptera. We focused on an atypical OR, SlitOR5, which exhibited a strong male-biased expression in antennae of the noctuid moth *S. littoralis* but did not group with the PR clade. We demonstrated, using a combination of heterologous expression and loss-of-function studies, that this OR is responsible for the male antennal response to (*Z,E*)-9,11-14:OAc, the major component of the *S. littoralis* sex pheromone blend. Due to the unexpected phylogenetic position of SlitOR5 outside of the previously defined PR clade, the question arose whether SlitOR5 is an exception or belongs to a previously unknown clade of moth PRs. This latter hypothesis is supported by the observation that the paralogous lineage containing SlitOR5 harbored many other sex-biased ORs, identified in species from six distinct lepidopteran families. Notably, male-biased ORs have been found in Lasiocampidae, Sphingidae, Noctuidae and Tortricidae. Among these, two ORs from *Ctenopseustis obliquana* and *C. herana* (Tortricidae) have been functionally studied by heterologous expression in cell cultures, but no ligand could be identified (Steinwender et al., 2015). In the Lasiocampidae species *Dendrolimus punctatus*, these male-biased ORs have been referred to as “Dendrolimus-specific odorant receptors”, with the suspicion that they would represent good PR candidates since in *Dendrolimus* species, there is no OR clustering in the PR clade (Zhang et al., 2014, 2018). No functional data yet confirmed this suspicion.

Conversely, within the SlitOR5 lineage, almost half of the sex-biased ORs were female-biased (13 out of 28, compared to 4 out of 63 in the classical PR clade). Female-biased ORs have been generally proposed to be involved in the detection of plant-emitted oviposition cues, as demonstrated in *Bombyx mori* (Anderson et al., 2009). However, another interesting hypothesis is that they could be tuned to male sex pheromones. In moths, little attention has been put on male pheromones, which are known to be involved in various mating behaviors such as female attraction, female acceptance, aggregation of males to form leks, mate assessment or inhibition of other males (reviewed in Conner and Iyengar, 2016). The use of male pheromone systems has been selected multiple times in distinct moth families, as reflected by the chemical diversity of male pheromone compounds and of the disseminating structures (Birch et al., 1990; Phelan, 1997; Conner and Iyengar, 2016). It is thus expected that this polyphyletic nature of male pheromones would result in a large diversity of female PR types. Accordingly, female-biased ORs were found in different clades within the phylogeny. However, most remain orphan ORs, including BmorOR30 that does belong to the SlitOR5 lineage but for which no ligand could be identified (Anderson et al., 2009). Although the most common male-emitted volatiles are plant-derived pheromones (Conner and Iyengar, 2016), some male courtship pheromones are long-chained hydrocarbons related to type I female pheromone compounds (Hillier and Vickers, 2004) that could be detected by female-biased type I PRs such as those identified within the SlitOR5 lineage.

The ancestral protein from which the so-called “PR clade” would have arisen is thought to be an OR tuned to plant-emitted volatiles (Yuvaraj et al., 2017, 2018). Here, we evidence that SlitOR5 is a new type I PR that belongs to a distinct early diverging lineage for which a role in pheromone detection had never been demonstrated. Together with the findings that PRs for Type 0 (Yuvaraj et al., 2017) and one PR for type II pheromones (Li et al., 2017) group in distinct paralogous lineages also unrelated to the PR clade, our data suggest that lepidopteran PRs have evolved four times in four paralogous lineages. Whether the SlitOR5 lineage has evolved from ORs that detected structurally-related plant volatiles - as has been proposed for type 0 (Yuvaraj et al., 2017) and classical type I PRs - remains elusive. Yet, no OR tuned to plant volatiles has been identified in closely related lineages. More functional data on SlitOR5 paralogs and orthologs in different moth species, possibly revealing more type I PRs, will help in understanding evolutionary history of this lineage, as to when and how these receptors have evolved, and confirm that we are facing a novel type I PR clade.

## Materials and methods

### Animal rearing and chemicals

*S. littoralis* were reared in the laboratory on a semi-artificial diet (Poitout and Buès, 1974) at 22°C, 60% relative humidity and under a 16h light:8h dark cycle. Males and females were sexed as pupae and further reared separately. *D. melanogaster* lines were reared on standard cornmeal-yeast-agar medium and kept in a climate- and light-controlled environment (25°C, 12h light:12h dark cycle). The 26 pheromone compounds used for electrophysiology experiments (Supplementary File 1) were either synthesized in the lab or purchased from Sigma-Aldrich (St Louis, MO, USA) and Pherobank (Wijk bij Duurstede, The Netherlands). Paraffin oil was purchased from Sigma-Aldrich and hexane from Carlo Erba Reagents (Val de Reuil, France).

### Quantitative real-time PCR

Total RNA from three biological replicates of 15 pairs of antennae of 2-day-old male and female *S. littoralis* was extracted using RNeasy Micro Kit (Qiagen, Hilden, Germany), which included a DNase treatment. cDNA was synthesized from total RNA (1 μg) using Invitrogen™ SuperScript™ II reverse transcriptase (Thermo Fisher Scientific, Waltham, MA, USA). Gene-specific primers were designed for *SlitOr5* (*Or5up*: 5’-TCGGGAGAAACTGAAGGACGTTGT-3’, *Or5do*: 5’-GCACGGAACCGCACTTATCACTAT-3’) and for the reference gene *SlitRpl13* (*Rpl13up*: 5’-GTACCTGCCGCTCTCCGTGT-3’, *Rpl13do*: 5’-CTGCGGTGAATGGTGCTGTC-3’). qPCR mix was prepared in a total volume of 10 μL with 5 μL of LightCycler^®^ 480 SYBR Green I Master (Roche, Basel, Switzerland), 4 μL of diluted cDNA (or water for the negative control) and 0.5 μM of each primer. qPCR assays were performed using the LightCycler^®^ 480 Real-Time PCR system (Roche). All reactions were performed in triplicate for the three biological replicates. The PCR program began with a cycle at 95°C for 13.5 min, followed by 50 cycles of 10 s at 95°C, 15 s at 60°C and 15 s at 72°C. Dissociation curves of the amplified products were performed by gradual heating from 55°C to 95°C at 0.5°C.s^−1^. A negative control (mix without cDNA) and a fivefold dilution series protocol of pooled cDNAs (from all conditions) were included. The fivefold dilution series were used to construct relative standard curves to determine the PCR efficiencies used for further quantification analyses. Data were analyzed with the LightCycler^®^ 480 software (Roche). Normalized expression of the *SlitOr5* gene was calculated with the Q-Gene software (Joehanes and Nelson, 2008) using *SlitRpl13* as a reference, considering it displays consistent expression as previously described in (Durand et al., 2010).

### Heterologous expression of SlitOR5 in Drosophila

The *SlitOr5* full-length open reading frame (1191 bp, GenBank acc. num. MK614705) was subcloned into the pUAST.attB vector. Transformant *UAS-SlitOr5* balanced fly lines were generated by BestGene Inc. (Chino Hills, CA, USA), by injecting the pUAST.attB-*SlitOR5* plasmid (Endofree prepared, Qiagen) into fly embryos with the genotype *y*^1^ *M{vas-int.Dm}ZH-2A w*\*; *M{3xP3-RFP.attP}ZH-51C* (Bischof et al., 2007), leading to a non-random insertion of the *UAS-SlitOr5* construct into the locus 51C of the second chromosome. The *UAS-SlitOr5* balanced line was then crossed to the *Or67d*^GAL4^ line (Kurtovic et al., 2007) to obtain double homozygous flies (genotype *w*; *UAS-SlitOr5,w*^+^; *Or67d*^GAL4^) expressing the *SlitOr5* transgene in at1 OSNs instead of the endogenous *Drosophila* receptor gene *Or67d*. The correct expression of *SlitOr5* was confirmed by RT-PCR on total RNA extracted from 100 pairs of antennae.

### Single-sensillum recordings

Single-sensillum extracellular recordings on *Drosophila* at1 OSNs were performed as previously described (de Fouchier et al., 2015). OSNs were stimulated during 500 ms, using stimulus cartridges containing 10 μg of pheromone (1 μg/μl in hexane) dropped onto a filter paper. Single-sensillum recordings on *S. littoralis* LT1A OSNs were performed using the tip-recording technique, as previously described (Pezier et al., 2007). Briefly, the tips of a few LT1 sensilla were cut off using sharpened forceps and a recording glass electrode filled with a saline solution (170 mM KCl, 25 mM NaCl, 3 mM MgCl_2_, 6 mM CaCl_2_, 10 mM HEPES and 7.5 mM glucose, pH 6.5) was slipped over the end of a cut LT1 sensillum. OSNs were stimulated with an air pulse of 200 ms (10 L.h^−1^), odorized using a stimulus cartridge containing 1 μg of pheromone (diluted at 1 μg/μl in hexane). Odorants were considered as active if the response they elicited was statistically different from the response elicited by the solvent alone (one-way ANOVA followed by a Tukey’s *post hoc* test).

### Heterologous expression of SlitOR5 in Xenopus oocytes and 2-electrode voltage-clamp recordings

Open reading frames of *SlitOr5* and *SlitOrco* (GenBank acc. num. EF395366, Malpel et al., 2008) were subcloned into the pCS2+ vector. Template plasmids were fully linearized with PteI for pCS2+-*SlitOR5* and NotI for pCS2+-*SlitOrco* and capped cRNAs were transcribed using SP6 RNA polymerase. Purified cRNAs were re-suspended in nuclease-free water at a concentration of 2 μg/μL and stored at −80°C. Mature healthy oocytes were treated with 2 mg/ml collagenase type I in washing buffer (96 mM NaCl, 2 mM KCl, 5 mM MgCl_2_ and 5 mM HEPES, pH 7.6) for 1–2 h at room temperature. Oocytes were later microinjected with 27.6 ng of *SlitOr5* cRNA and 27.6 ng of *SlitOrco* cRNA. After 4 days of incubation at 18°C in 1× Ringer’s solution (96 mM NaCl, 2 mM KCl, 5 mM MgCl2, 0.8 mM CaCl2, and 5 mM HEPES, pH 7.6) supplemented with 5% dialyzed horse serum, 50 mg/ml tetracycline, 100 mg/ml streptomycin and 550 mg/ml sodium pyruvate, the whole-cell currents were recorded from the injected oocytes with a 2-electrode voltage clamp. Oocytes were exposed to the 26 pheromone compounds diluted at 10^−4^ M in Ringer’s solution, with an interval between exposures which allowed the current to return to baseline. Data acquisition and analysis were carried out with Digidata 1440A and pCLAMP™10 software (Molecular Devices, San Jose, CA, USA). Odorants were considered as active if the mean response they elicited was statistically different from 0 (Wilcoxon signed rank test). Dose–response experiments were performed using pheromone concentrations ranging from 10^−9^ up to 10^−5^ M and data were analyzed using GraphPad Prism 5.

### Phylogenetic analyses

The dataset of lepidopteran amino acid OR sequences used to build the phylogeny shown in Fig. 1 included entire OR repertoires from the following 9 species, each belonging to a different lepidopteran superfamily: *Bombyx mori* (Bombycoidea; Tanaka et al., 2009), *Dendrolimus punctatus* (Lasiocampoidea; Zhang et al., 2018), *Ectropis grisescens* (Geometroidea; Li et al., 2017), *Epiphyas postvittana* (Tortricoidea; Corcoran et al., 2015), *Eriocrania semipurpurella* (Eriocranioidea; Yuvaraj et al., 2017), *Heliconius melpomene* (Papilionoidea; The Heliconius Genome Consortium, 2012), *Ostrinia furnacalis* (Pyraloidea; Yang et al., 2015), *Plutella xylostella* (Yponomeutoidea; Engsontia et al., 2014) and *Spodoptera littoralis* (Noctuoidea; Walker et al., 2019). The dataset used to build the phylogeny shown in Fig. 4 contained amino acid sequences from the same 9 species falling into that clade, plus all the sequences of ORs within that clade showing a marked sex-biased expression (threshold of a 10-fold difference in expression rate between male and female antennae, based on RNAseq or qPCR experiments) and/or for which ligands have been identified as of September 2018 (Supplementary File 2). Alignments were performed using Muscle (Edgar, 2004) as implemented in Jalview v2.10.5 (Waterhouse et al., 2009). Phylogenetic reconstruction was performed using maximum likelihood. The best-fit model of protein evolution was selected by SMS (Lefort et al., 2017) and tree reconstruction was performed using PhyML 3.0 (Guindon et al., 2010). Node support was first assessed using 100 bootstrap iterations, then the file containing bootstrap trees was uploaded on the Booster website (Lemoine et al., 2018) to obtain the TBE (Transfer Bootstrap Expectation) node support estimation. Figures were created using the iTOL web server (Letunic and Bork, 2016).

### SlitOr5 knock-out via CRISPR/Cas9 and mating behavior assessment

A guide RNA (gRNA sequence: AGCATAAATACTGGACCCAG TGG) was designed against the first exon of the *SlitOr5* gene using the CRISPOR gRNA design tool (crispor.tefor.net, Haeussler et al., 2016) and transcribed after subcloning into the DR274 vector using the HiScribe™ T7 High Yield RNA Synthesis Kit (New England Biolabs, Ipswich, MA, USA). The Cas9 protein was produced in *Escherichia coli* as previously described (Ménoret et al., 2015). A mix of Cas9 protein and gRNA was injected in freshly laid eggs using an Eppendorf - Transjector 5246, as previously described (Koutroumpa et al., 2016). Individual genotyping at every generation was performed via PCR on genomic DNA extracted from larvae pseudopods (Wizard^®^ Genomic DNA Purification Kit, Promega, Madison, WI, USA) using gene-specific primers (forward: 5’-CCAAAAGGACTTGGACTTTGAA-3’; reverse: 5’-CCCGAATCTTTTCAGGATTAGAA-3’) amplifying a fragment of 728 bp encompassing the target sequence. Mutagenic events were detected by sequencing the amplification products (Biofidal, Vaulx-en-Velin, France). G0 larvae carrying a single mutagenic event were reared until adults and crossed with wild-type individuals. Homozygous G2 individuals were obtained by crossing G1 heterozygous males and females. Mating capacity of homozygous mutants were measured by placing a wild-type virgin mature individual together with a same-age-mutant from the opposite sex (*n* = 10 for each males and females) in a plastic box at the beginning of the third scotophase in the rearing chamber. Couples were observed under dim red light every hour for 15 hours. If no mating occurred, the boxes were kept in the same conditions for several days until adult death and egg batches, if any, were counted every day.

### Phenotyping of CRISPR/Cas9 mutants by electroantennogram recordings

Electroantennogram recordings were performed as previously described (Koutroumpa et al., 2016) on isolated male antennae from wild-type animals and from heterozygous and homozygous *SlitOr5* mutants. Antennae were stimulated using stimulus cartridges loaded with 10 μg of linalool or benzyl alcohol diluted in paraffin oil, and 1 μg of (*Z,E*)-9,11-14:OAc or (*Z,E*)-9,12-14:OAc diluted in hexane. Stimulations lasted for 500 ms (30 L/h). Negative controls consisted of paraffin oil and hexane alone. The maximum depolarization amplitude was measured using the pCLAMP™10 software. Normalized mean responses were calculated (response to the solvent was subtracted) and data were analyzed using a one-way ANOVA followed by a Tukey’s *post hoc* test.

## Acknowledgements

We are grateful to Lixiao Du and Françoise Bozzolan for technical assistance, Bin Yang and Thomas Chertemps for their help and advice, and Pascal Roskam and Philippe Touton for insect rearing. We also thank Christophe Héligon (CRB Xénope, Rennes) for providing the pCS2+ plasmid. This work has been funded by the French National Research Agency (ANR-16-CE02-0003-01 and ANR-16-CE21-0002-01 grants), the National Natural Science Foundation of China (31725023, 31621064) and a PRC NSFC-CNRS 2019 grant.

## Competing interests

The intellectual property rights of SlitOR5 have been licensed by Inra, Sorbonne Université and CNRS for the purpose of developing novel insect control agents.

## Source data files

Figure 1 – **source data 1.** Mean Normalized Expression values of *SlitOr5* measured in the 3 biological replicates.

Figure 1 – **source data 2.** Alignment of amino acid sequences used to build the phylogeny (FASTA format).

Figure 2 – **source data 1.** Raw results of electrophysiology experiments.

Figure 3 – **source data 1.** Raw results of the EAG experiment.

Figure 4 – **source data 1.** Alignment of amino acid sequences used to build the phylogeny (FASTA format).

## Supplementary files

**Supplementary File 1.** List of synthetic compounds used for electroantennogram and single-sensillum recordings.

**Supplementary File 2.** Functional and sex-biased expression data available for lepidopteran pheromone receptors (as of September 2018).

